# Plant organ evolution revealed by phylotranscriptomics in *Arabidopsis*

**DOI:** 10.1101/119511

**Authors:** Li Lei, Joshua G Steffen, Edward J Osborne, Christopher Toomajian

## Abstract

The evolution of species’ phenotypes occurs through changes both in protein sequence and gene expression levels. Though much of plant morphological evolution can be explained by changes in gene expression, examining its evolution has challenges. To gain a new perspective on organ evolution in plants, we applied a phylotranscriptomics approach. We combined a phylostratigraphic approach with gene expression based on the strand-specific RNA-seq data from seedling, floral bud, and root of 19 *Arabidopsis thaliana* accessions to examine the age and sequence divergence of transcriptomes from these organs and how they adapted over time. Our results indicate that, among the sense and antisense transcriptomes of these organs, the sense transcriptomes of seedlings are the evolutionarily oldest across all accessions and are the most conserved in amino acid sequence for most accessions. In contrast, among the sense transcriptomes from these same organs, those from floral bud are evolutionarily youngest and least conserved in sequence for most accessions. Different organs have adaptive peaks at different stages in their evolutionary history, however, from the Magnoliophyta stage to the Brassicale stage, all three organs show a common adaptive signal. Our research is significant because it offers novel evolutionary insight on plant organs revealed by phylotranscriptomics.

## INTRODUCTION

The evolution of species’ phenotypes occurs at two levels, not only through changes in protein sequence but also in gene expression levels (*i.e*., changes in the transcriptome)^1^. Though early molecular evolutionary studies focused on protein evolution, data from subsequent research supported the extremely important role gene expression changes played in phenotypic evolution in both animals ^2
^ and plants ^3^. However, examining the evolution of gene expression in different organs presents novel challenges compared to examining the evolution of protein coding sequence. Recently, Yang and Wang, through comparing the transcriptomes in different tissues of maize, rice and *Arabidopsis*, found a correlation across genes in the rates of sequence evolution and divergence in gene expression patterns; and also evidence that the expression differences of the orthologous genes among different species varied between different organs ^4^. Nevertheless, the age and sequence divergence level of those transcriptomes in different organs and how those organs adapted in different stages along their evolutionary history are unknown.

Transcriptome profiling by RNA-sequencing facilitates comparisons of gene expression across organs in the context of coding sequence evolution. Comparisons of gene expression in different organs combined with their coding sequence evolution will illuminate how these processes jointly shape the phenotypic diversity of the animal and plant kingdoms.

Due to DNA’s double-stranded complementarity, the mRNA transcripts that are used as templates for protein translation look like what is referred to as the sense strand of the DNA with respect to the protein-coding gene, and the whole process called sense expression. However, certain genes display measurable antisense expression, in which mRNAs are transcribed in the opposite (antisense) direction and thus contain sequence complementary to the sense mRNA. Natural antisense transcripts (NAT) produced by antisense expression can regulate the transcript abundance of their complements by triggering the biogenesis of natural antisense short interfering RNAs (nat-siRNAs) that subsequently guide transcript cleavage ^5-7^. Additionally, antisense expression can have a function in antisense transcript-induced RNA splicing, alternative splicing, and polyadenylation, which can regulate gene expression abundance and protein-coding complexity ^8^.

A novel approach for comparing the gene expression in different organs in conjunction with sequence evolution is to estimate the contribution of genes from different “phylostrata” ^9,10^ to gene expression. The evolutionary origin of any gene can be traced by sequence similarity searches in genomes representing the whole tree of life, an approach known as “phylostratigraphy”. In this approach, every gene within a genome has a phylogenetic rank, and is associated with a “phylostratum” ^10^ based on its inferred phylogenetic emergence. By combining phylogenetic hierarchy and gene expression to examine the developmental hourglass model, which predicts the pattern of morphological divergence for different developmental stages, in zebrafish, Domazet-Loso *et al*. first introduced the transcriptome age index (TAI), which integrates the age of a gene with its expression level at a given developmental stage and sums this over all genes expressed at the respective stage ^10^. Similarly, Quint *et al*. introduced the transcriptome divergence index (TDI), which integrates the sequence divergence of a gene with its expression level at a given developmental stage and sums this over all genes expressed at the respective stage, to explore the embryonic developmental hourglass in *Arabidopsis* ^11^. More recently, using TAI and TDI, Cheng *et al*. investigated the developmental hourglass in fungi ^12^. Although the studies mentioned above applied the indices TAI and TDI, their application was only limited to the examination of the developmental hourglasses in different organisms. No studies have used these two indices to investigate organ evolution, or more specifically to examine the age and sequence divergence of transcriptomes in different organs, either in animals, plants, or microorganisms.

It is well known that root, flower, stem and leaves (the latter two are major components of seedlings) are important organs for the majority of plants. Thus, knowing more about the evolutionary properties of their transcriptomes, and how those organs adapted in different stages of evolutionary history could be of great significance in studying a plant organ’s origin or development, and even for improving a plant’s adaptation to the environment. Nevertheless, how and when these organs evolved, is not well understood, although there are estimates of those organs’ origins. Some of the earliest fossil evidence of roots comes from 419-408 million years ago (mya) in clubmosses (*e.g*., *Drepanophycus spinaeformis*) and their close relatives in the extinct zosterophylls (*e.g*., *Bathurstia denticulate*) ^13^. Flower-like structures first appeared in the fossil record ∼130 mya in the Cretaceous ^14^. Leaves first evolved during the Late Devonian to Early Carboniferous, diversifying rapidly until the designs settled down in the mid Carboniferous ^15^. Although one can infer the origin of each organ of plants from the fossil record, details of the adaptive history for each organ are still lacking, specifically, in which phylostrata are the organ-specific expressed genes enriched (indicating an adaptive stage for plant organs). Several pioneer studies can serve as models for examining the adaptive profiles of plant organs because they have used the approach of mapping genes specifically expressed from domains in the vertebrate head sensory system and brain to their phylostrata ^16,17^ in order to reveal the adaptive history of each domain.

Here we examine the age and sequence divergence of the transcriptomes from different organs in *Arabidopsis* to reveal the adaptive profile of each organ along its evolutionary history. We applied the phylostratigraphic approach and combined it with gene expression based on the strand-specific RNA-seq data from seedling, floral bud, and root of 19 *Arabidopsis thaliana* accessions. Our results indicate that, among the sense and antisense transcriptomes of these organs, the sense transcriptomes of seedlings are the evolutionarily oldest across all accessions and are the most conserved in amino acid sequence for most accessions. In contrast, among the sense transcriptomes from these same organs those from floral bud are evolutionarily youngest and least conserved in sequence for the majority of accessions.

## RESULTS

### Analyses of gene expression in *Arabidopsis* seedling, root and floral bud

We collected the strand-specific transcriptomes from three organs: root, seedling and stage 12 floral buds^18^ (flower for short hereafter) from 19 *Arabidopsis* accessions, which are the founders of the MAGIC lines ^19,20^. Then we normalized the expression as reads per million (RPM). Due to the noise inherent in estimating gene expression by RNA-seq and the potential for polymorphism across the 19 accessions, we classified a gene as expressed in a certain organ (separately for both sense and antisense expression) if the expression abundance in at least 5 accessions was above 0.8 RPM in that organ. Figures 1A (sense expression) & 1B (antisense expression) indicate the expression breadth of all expressed genes. Given our three organs, we defined seven expression groups: specific to root (R), flower (F), and seedling (S); shared between root-seedling (RS), flower-root (FR), and seedling-flower (RS), and shared among root-seedling-flower (RSF). More than half of sense expressed (85.63%) and antisense expressed (55.46%) transcripts are expressed in more than a single organ (Figure 1A & B). Organ-specific expressed genes and genes expressed in only two organs are much more frequent for antisense compared with sense expression, while the genes shared by three organs are underrepresented in antisense expression (Figure 1A & B; Table S1).

**Figure 1.**
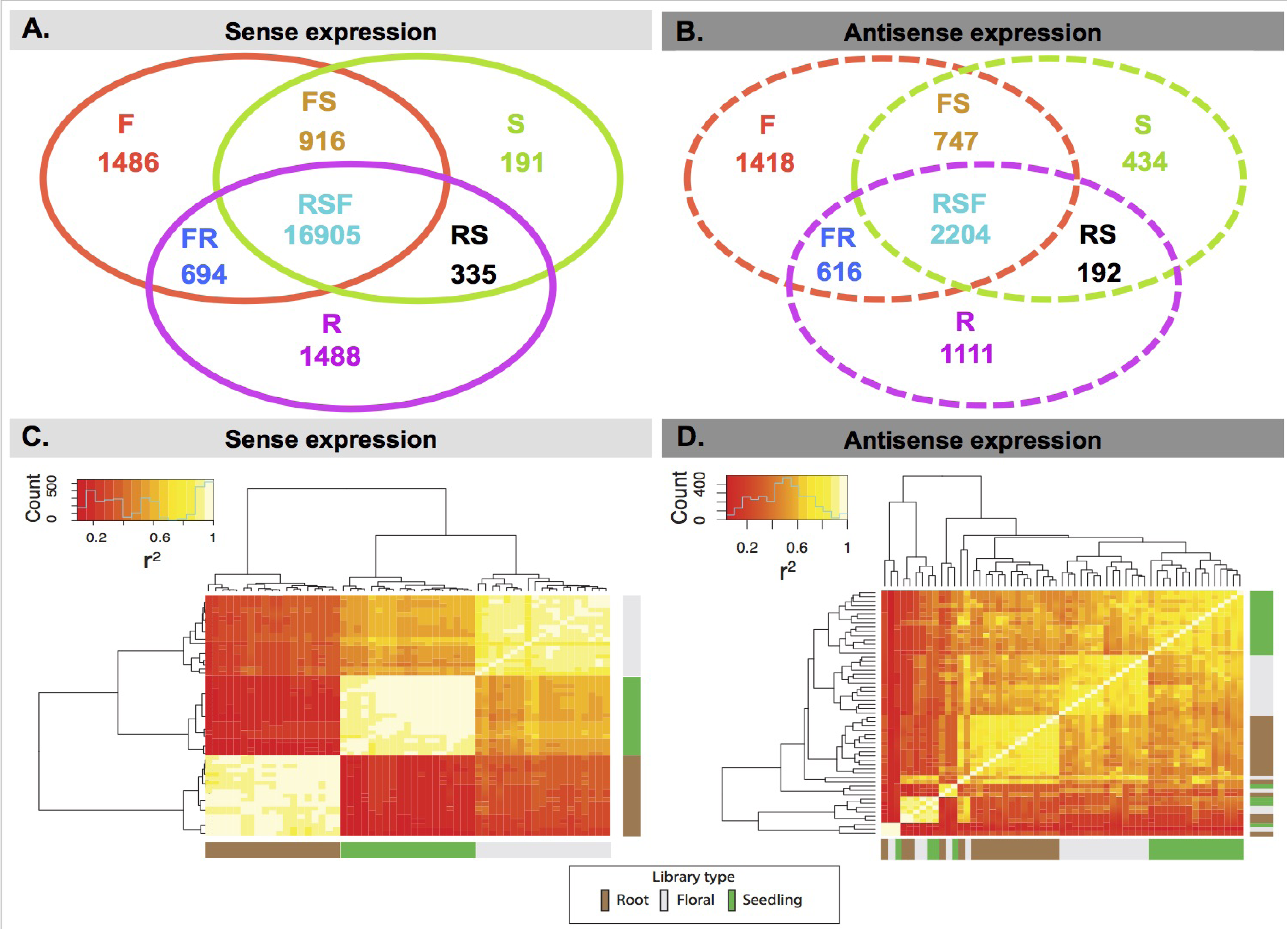
Gene expression in root, flower and seedling across 19 *A. thaliana* accessions. A. Venn diagram of the counts of genes with sense expression in root, flower and seedling across 19 accessions. B. Venn diagram of the counts of genes with antisense expression in root, flower and seedling across 19 accessions. C. Heatmap of sense expression in three organs across 19 accessions. D. Heatmap of antisense expression in three organs across 19 accessions. F: Flower-specific expressed genes; S: Seedling-specific expressed genes; R: Root-specific expressed genes; FS: genes with expression shared by Flower and Seedling only; FR: genes with expression shared by Flower and Root only; RS: genes with expression shared by Root and Seedling only; RSF: genes with expression shared by Root, Seedling and Flower.

Additionally, we clustered each organ from each accession based on gene expression (Figure 1C & D). The results showed that sense expression between accessions within organs is highly correlated while antisense expression is more variable across accessions.

### Phylostrata and dN/dS ratio

Any gene can be mapped to the time point—the phylostratum—when its oldest domain emerged in evolution ^10^. Following the approach of Drost, *et al*.*^21^*, the 25,260 protein-coding genes of *A. thaliana* (TAIR 10) were assigned into 12 phylostrata (Figure 2A). The phylostrata of those genes cover a time span from the origin of cellular organisms (ca. 2520 Ma) to the terminal lineage (0 Ma), that is, *A. thaliana*. More than half of genes (64.87%) originated from the three most ancient phylostrata (Cellular organisms, Eukaryota, and Viridiplantae), which indicates that relatively few genes originated during the subsequent evolutionary processes along the land plant specific portion of the lineage leading to *A. thaliana*. Nevertheless, these later appearing genes may have played an extremely important role in the development and divergence of seed plants. For the rest of the phylostrata, the largest set of genes appeared in Embryophyta (10.43%), then *A. thaliana* (6.94%), and then Magnoliophyta (6.22%). That could indicate that those three phylostrata could be key phylostrata in their ultimate contribution to *A. thaliana* evolution.

**Figure 2.**
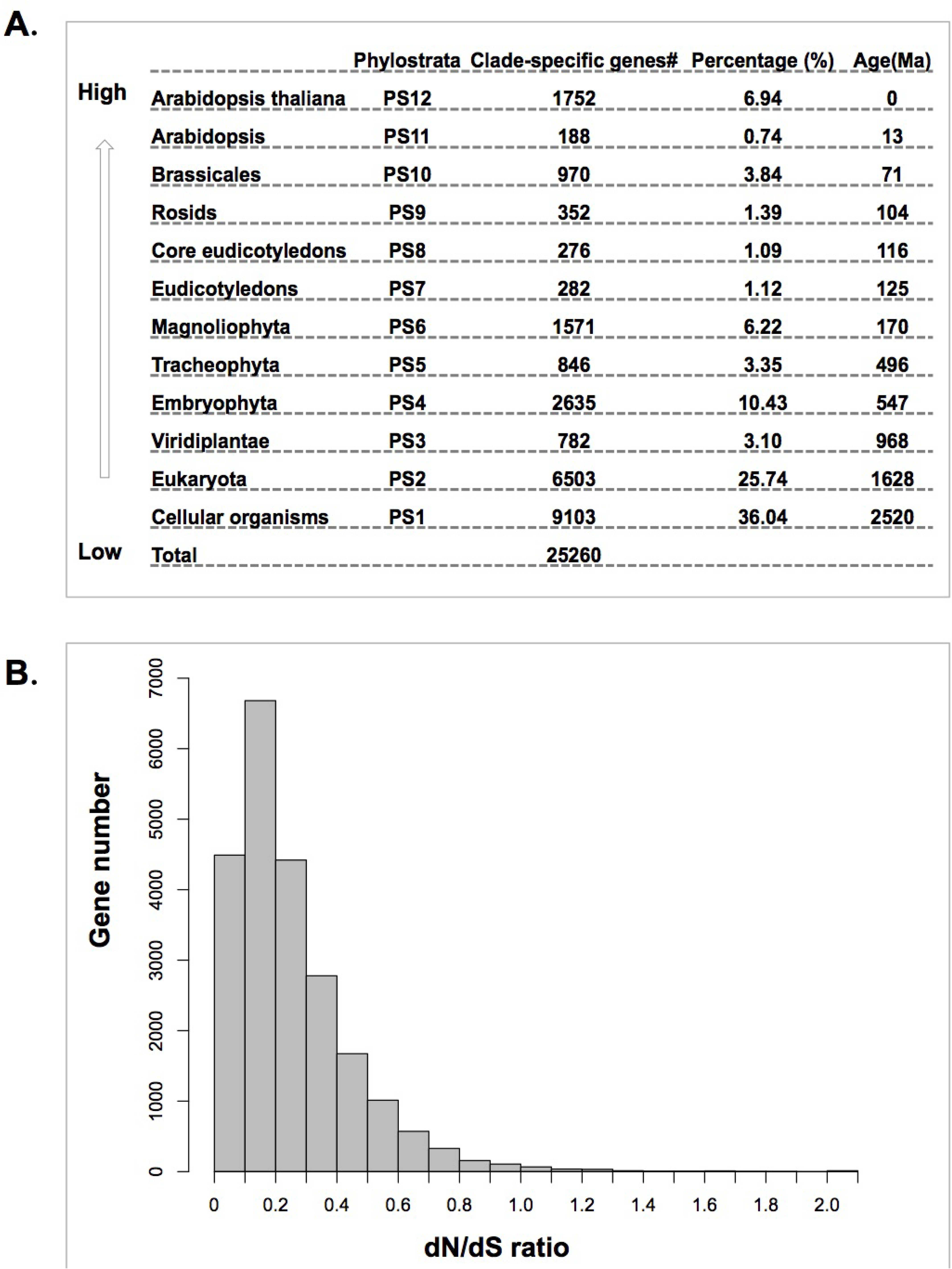
Phylostrata and *dN/dS* ratio for protein coding genes in *A. thaliana*. A. Phylostrata and the number of genes mapped to each phylostratum. B. The distribution of Nonsynonymous substitution rate (dN)/Synonymous substitution rate (*dS*) for orthologous gene pairs of *A. thaliana* and *A. lyrata*.

The dN/dS ratio is used to assess the selective pressure on protein-coding regions and reflects natural selection. The majority of the orthologous gene pairs between *A. thaliana* and other species in Brassicaceae have a dN/dS ratio less than 1 (Figure 2B and Figure S1), indicating purifying selection predominates in the majority of gene pairs.

## Evolutionary age of *Arabidopsis* seedling, root and flower transcriptomes

In order to estimate the age of each transcriptome, we computed a mean TAI and standard error based on phylostratigraphy for each organ of each accession with 1000 bootstrap replicates. The TAI quantifies the mean evolutionary age of the transcriptome in a certain organ; the lower the TAI, the evolutionarily older the transcriptome ^9,11^. For sense expression across 19 accessions, the evolutionarily older genes tended to be expressed in seedling (consistently lowest average TAI across all accessions), while the younger genes tended to be expressed in flower (highest average TAI in all but three accessions, Wil-2, Zu-0 and Bur-0, in which root has the highest average TAI rather than flower) (Figure 3A and Table S2-3). This result is also clearly evident in the comparison of relative expression between old genes (PS1-PS3) and young genes (PS4-PS13) (Figure 3E and Figure S3). It shows that old genes have much higher relative sense expression in seedling than young genes; in contrast, in root and flower, young genes have relatively higher sense expression than old genes. In particular, the magnitude of the difference between young genes and old genes tends to be bigger in flower than in root (Figure 3E and Figure S3). Additionally, the pattern of TAI is fairly consistent across accessions, with the minor exception of three accessions in which root has the highest average TAI rather than flower.

**Figure 3.**
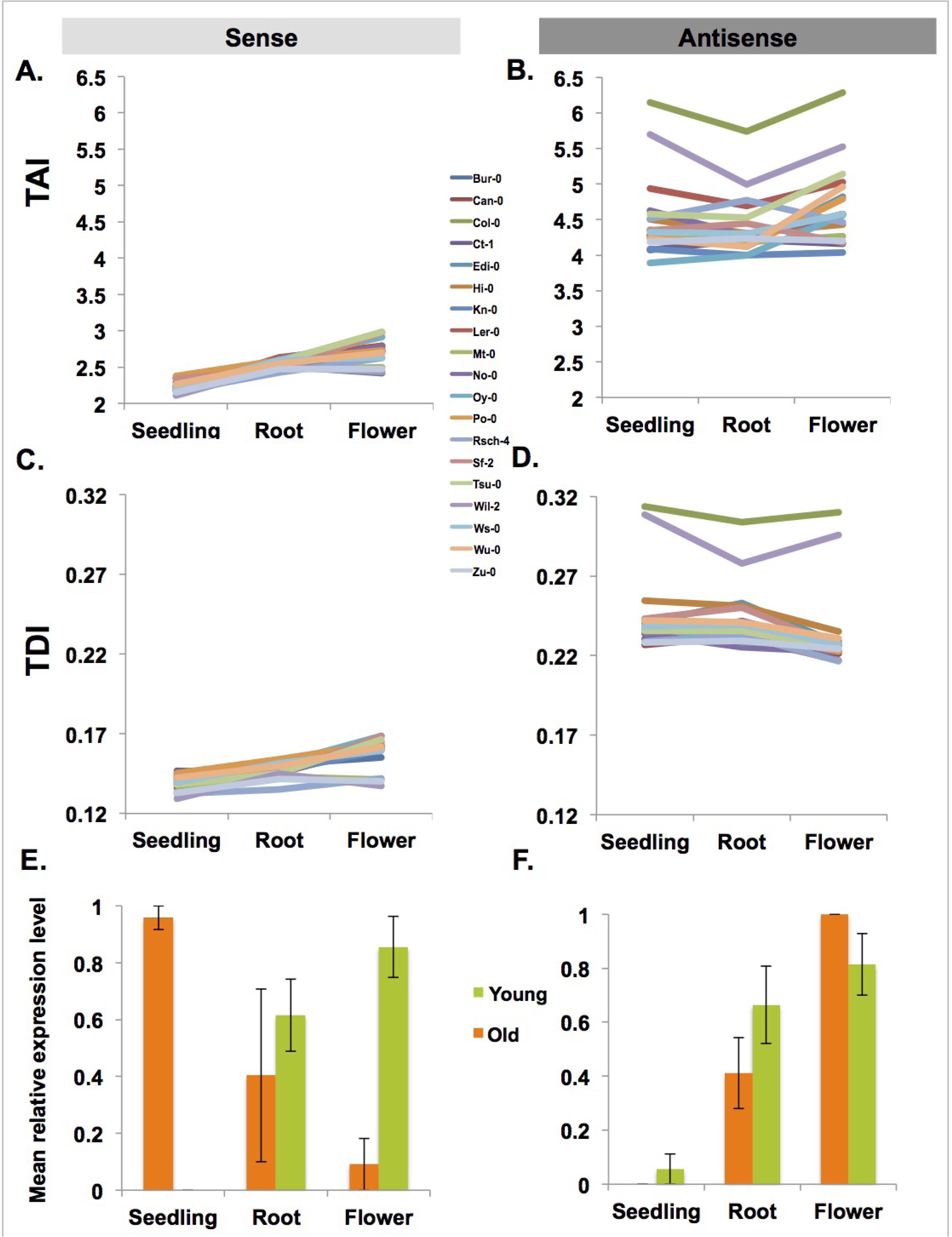
Transcriptome indices of seedling, root and flower across 19 *A. thaliana* accessions. A. The sense transcriptome age index (TAI) profile of three organs across 19 accessions. B. The antisense transcriptome age index (TAI) profile of three organs across 19 accessions. C. The sense transcriptome divergence index (TDI) profile of three organs across 19 accessions. D. The antisense transcriptome divergence index (TDI) profile of three organs across 19 accessions. The standard errors of TAI and TDI, estimated by bootstrap analysis (1000 replicates), are extremely small so that they cannot be seen in A-D, but are presented in Tables S2, 4, 6, 8 & 10. The comparisons of TAI and TDI between any two organs are significant by the Mann Whitney test, and the p-values are presented in Tables S3, 5, 7, 9 & 11. E. Mean relative sense expression levels of evolutionarily old (PS1-PS3) and young (PS4-PS12) genes in different organs in a representative accession: Ler-0. F. Mean relative antisense expression levels of evolutionarily old (PS1-PS3) and young (PS4-PS13) genes in different organs in Ler-0.

In contrast, for antisense expression there are no major consistent differences in TAI across organs (Figure 3B & F; Figure S3). Instead, two accessions (Col-0 and Wil-2) have a much higher value for TAI (Figure 3B and Table S2). That could be due to a larger number of young genes that show antisense expression in a restricted set of accessions, including Col-0 and/or Wil-2. A bias in the analysis of accession-specific genes, which also must correspond to the *A. thaliana* phylostratum (PS12) if they resulted from gene duplication or creation, predicts this result for Col-0. Specifically, though accession-specific genes are expected in all of the accessions in the study, using Col-0 as the reference to define the set of genes for the phylostrata analysis (as we have done) can create a bias in TAI calculations across the set of accessions, because genes specific to other accessions but absent in Col-0 are not included in the analyses. On the other hand, genes specific to Col-0 are included in the TAI calculation but must not be expressed in other accessions. As a consequence Col-0 and the accessions most closely related to it are expected to have slightly higher average expression from the class of the youngest genes relative to other accessions. This would tend to increase TAI for Col-0, which is what we see, at least for the TAI calculation for antisense expression.

Compared to the sense expression of genes, the TAI of antisense expression is much higher for all three organs (Figure 3A & B), which indicates that antisense expression tends to occur in younger genes.

Moreover, the standard errors of TAI in both sense and antisense across 19 accessions are fairly small (Figure 3A & B and Table S2).

## Sequence divergence of *Arabidopsis* seedling, root and flower transcriptomes

We computed a transcriptome divergence index (TDI) based on dN/dS ratios for each organ. TDI quantifies the mean selection force acting on the coding sequence of a transcriptome; the further below 1 the TDI, the stronger the force of purifying selection ^22^. We found that TDI exhibited a similar pattern to the TAI for both sense and antisense expression. For sense expression, seedling tended to have the lowest average TDI, giving the strongest signal of purifying selection and greatest amino acid conservation, and floral bud tended to have the highest average TDI, suggesting its transcriptome was experiencing weaker purifying selection and therefore greater amino acid divergence (Figure 3C; Figure S2 A, C & E, Table S4, 5, 6, 7, 8, 9, 10&11). The pattern of TDI for sense expression is relatively consistent across accessions, with only five accessions deviating from this general pattern when using TDI computed with dN/dS ratios estimated from comparing *A. thaliana* to any of four related species. The lowest average TDI value occurs in root rather than seedling for Ct-1 (across all four TDI calculations) and Sf-2 (in two of the four TDI calculations), while the highest average TDI value occurs in root rather than floral bud for Wil-2 (across all four TDI calculations), Zu-0 (in two of the four TDI calculations) and Mt-0 (in one of the four TDI calculations).

In contrast to the consistent pattern of TDI for sense expression across accessions, for antisense expression there is no consistent pattern of TDI with respect to organs and relatively higher variance in TDI across accessions (Figure 3D and Figure S2 B, D & F). Instead, two accessions (Col-0 and Wil-2) have a much higher value for TDI when using dN/dS ratios computed from comparisons of *Arabidopsis thaliana* with *A. lyrata* (Figure 3D, Table S4), but do not show similar elevated TDI values when using dN/dS ratios computed from a comparison of *A. thaliana* with *T. halophila* (Figure S2B and Table S6), *C. rubella* (Figure S2D and Table S8) or *B. rapa* (Figure S2F and Table S10). This could result from a reason similar to the bias in TAI due to accession-specific genes explained above. In this case, the relevant genes must be present in the closest related species (or else divergence could not be calculated), but nevertheless many genes can still be polymorphic for presence/absence among *A. thaliana* accessions. The bias occurs because genes are only included in the analysis when they are found in Col-0, but can be absent from other accessions. Assuming that presence/absence genes tend to have less selective constraint, and hence a higher dN/dS ratio, Col-0 is expected to have an inflated ratio. Once we make comparisons with relatives more distant than *A. lyrata*, the inflation in the Col-0 TDI greatly decreases or disappears. This would be expected if genes with presence/absence polymorphisms are largely limited to those that arose after the *A. thaliana* lineage split from all but its closest relative, so that the bias specific to Col-0 and the accessions most closely related to it disappears once the analysis is restricted to genes for which divergence from the more distant relatives can be estimated.

Compared to sense expression, the TDI of antisense expression is much higher across all three organs.

Similar to TAI, the standard errors of TDI in both sense and antisense expression across 19 accessions are also fairly small (Figure 3C & D and Tables S4, 6, 8 & 10).

## Adaptive patterns in *Arabidopsis* seedling, root and floral bud

To further shed light on the history of adaptation (estimated as the enrichment of the organ-specific expressed genes in different phylostrata) in seedling, root and flower, we mapped seedling-specific (S), root-specific (R) and flower-specific (F) expressed genes (Figure 1A and B) (for both sense and antisense expression) to their corresponding phylostrata, and plotted the adaptive profiles of these genes (Figure 4 and Figure S4). We also mapped the genes expressed in all three organs (RSF) (Figure 1A and B) to phylostrata, and plotted the adaptive profile for comparison (Figure 4 and Figure S4). Generally, different organs have different adaptive peaks (over-represented phylostrata) (Figure 4). The adaptive peaks for floral bud are at the Embryophyta, Magnoliophyta, Eudicotyledons, core Eudicotyledons, Rosids and Brassicales phylostrata, although only in Eudicotyledons and Brassicales is the over-representation significant. The first adaptive peak appears at Embryophyta, though the level of over-representation is low, suggesting that flower formation could partially be traced back to genes originating in Embryophyta ancestors. The second adaptive peak appears at the Magnoliophyta phylostratum. Flower-specific genes are also overrepresented in Eudicotyledons, core Eudicotyledons, Rosids and Brassicales, suggesting that novel flower-specific genes continued to originate at different stages of plant evolution after Magnoliophyta to sustain the complex character and activity of flowers. But after Brassicales there is no enrichment in *Arabidopsis* and *A. thaliana*, possibly indicating that new genes that have evolved since Brassicales are not often expressed in flowers.

**Figure 4.**
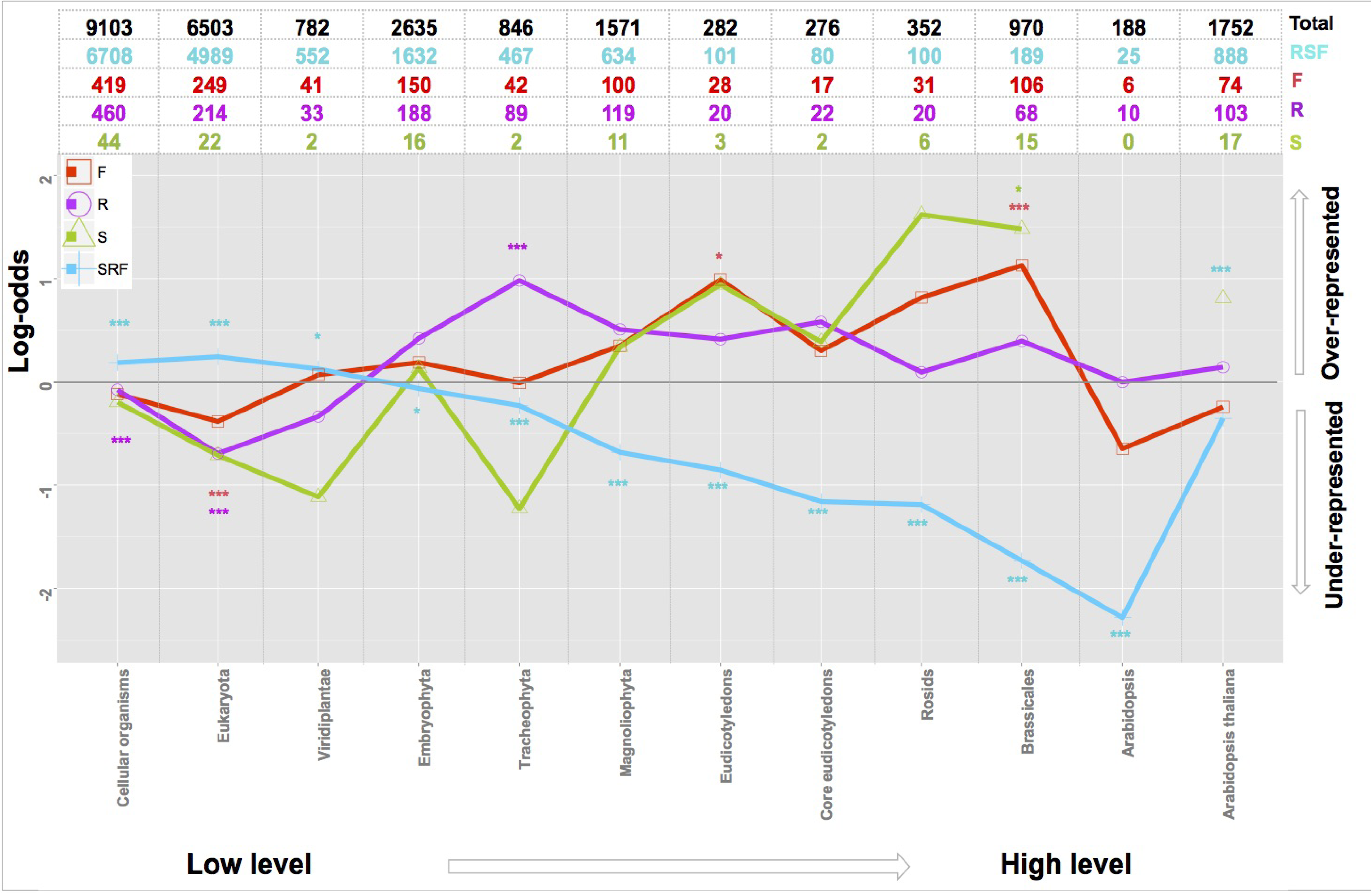
Enrichment analysis of organ-specific sense-expressed genes in different phylostrata. RSF: genes with expression shared by Root, Seedling and Flower; S: Seedling-specific expressed genes; R: Root-specific expressed genes; F: Flower-specific expressed genes; Total: all the protein coding genes with an assigned phylostratum. Gray line: A log-odds of zero, which corresponds to the actual number of organ-specific genes in each phylostratum equaling the predicted number. (* P <0.05, ** P <0.01 and *** P <0.001).

The adaptive peaks for root are at Embryophyta, Tracheophyta, Magnoliophyta, Eudicotyledons, core Eudicotyledons, Rosid, Brassicales and *A. thaliana* (Figure 4). The biggest adaptive peak for root is in Tracheophyta. This indicates that Tracheophyta is the most important phase during the evolutionary history of the organ, consistent with the fact that starting from Tracheophyta, plants have had independent roots, and that the earliest fossils of roots indicate they originated from early species within Tracheophyta ^23^.

The adaptive peaks for seedling are at Embryophyta, Magnoliophyta, Eudicotyledons, core Eudicotyledons, Rosids, Brassicales and *A. thaliana* (Figure 4). Additionally, we also observed that before Embryophyta, none of the previous phylostrata are overrepresented in the three sets of organ-specific sense-expressed genes, but the RSF shared genes show over-representation for each of the three early phylostrata (Figure 4), which could be because root, seedling and floral bud were not differentiated before this phylostratum, also indicating that few new genes with broad expression across all 3 organs have originated since root, seedling, and floral bud have differentiated.

Investigating the adaptive pattern of genes with antisense expression in different organs (Figure S4), we found that for all the groups, in all of the phylostrata before Magnoliophyta, genes with organ-specific antisense expression were distributed fairly evenly without strong over-representation (with log-odd less than 0.5) except in Viridiplantae. In phylostrata 11 (*Arabidopsis*), all groups of genes were strongly under-represented, which indicates that antisense expression of genes that originated in this phylostratum is very rare, and hence could not play a major role in regulating gene expression for these genes. The overrepresentation value only exceeds 0.5 in core Eudicots and Viridiplantae; and both are in root. This could indicate that during the evolution of root, antisense expression could have been important to regulate the gene expression in the phylostrata of Viridiplantae and core Eudicots. Also in Embryophyta, even though the log-odds are less than 0.5, the root and the RSF expressed genes show significant enrichment, which could suggest that antisense expression played an important role in regulating gene expression in Embryophyta.

## Gene ontology analysis for organ-specific genes and genes expressed in all three organs

Since we investigated the adaptive patterns for seedling, flower and root, we were curious about the functional categories to which organ-specific genes belong, focusing on sense expression specifically. We performed the Gene Ontology analysis for R, S and F genes considering all the genes expressed in at least one organ as background. In parallel, weperformed a GO analysis for genes in the RSF set (Table S14). We found that flower-specific genes tend to be enriched in several terms, including: pollen tube development, plant-type cell wall and cell morphogenesis related, and reproductive processes related (Table S12). Root-specific genes tend to be enriched in several terms, including: oxidoreductase related, transport related terms related to response to different biotic and abiotic factors, and root development related terms (Table S13). Genes expressed in three organs tend to be enriched in the following terms: organelle related processes, cytoplasmic related processes, cellular metabolite processes, etc. (Table S14). However, we found no GO terms enriched in seedling-specific genes, which could be because of the small number of seedling-specific sense-expressed genes.

## DISCUSSION

Plants play an important role in life on land, and the emergence of their morphological differences correlates with speciation, the ecological consequences of variation in physiological or developmental traits, and the adaptive evolution of different species ^24^. Developing an understanding of the evolutionary origins of different organ systems lies at the heart of evolutionary biology. A direct approach to examine the origins and evolution of plant organs uses fossils ^25,26^. The fossil record alone cannot comprehensively reveal the evolutionary processes for plant species ^26^. Here we took a novel approach, phylotranscriptomics. This indirect approach combines gene expression and sequence evolution to investigate the age and sequence divergence of the transcriptomes in different organs and the adaptive profile of those organs along their evolutionary history.

TAI reflects long-term evolutionary changes covering 4 billion years since the origin of life, and TDI reflects short-term evolutionary changes covering 5-16 million years since the divergence of *A. thaliana* and four other Brassicaceaes ^49,51-53^. The results of TAI and TDI analyses for sense expression indicate that the transcriptome in flower is youngest and has the highest amino acid sequence divergence, the root transcriptome is much older with much lower sequence divergence, and finally the transcriptome in seedling is oldest and most conserved in sequence, experiencing more purifying selection. This may be consistent with the theory of plant organ evolution and the earliest fossil records of plant organs. According to Hagemann’s theory, the most primitive land plants that gave rise to vascular plants were flat, thalloid, leaf-like, without axes, somewhat like a liverwort or fern prothallus. Axes such as stems and roots evolved later as new organs ^27
,28^. Roots evolved in the sporophytes of at least two distinct lineages of early vascular plants (Tracheophyta) during their initial major radiation on land in Early Devonian times (c. 410–395 million years ago) ^23^. Real flowers are modified leaves possessed only by Magnoliophyta, a group that originated and diversified during the Early Cretaceous ^29^. Flowers are relatively late to appear in the fossil record. However, there are some fossil records of flower-like organs in ferns or cycads and gnetales ^22^. Seedlings include the stems and leaves, both of which are more primitive than roots and flowers ^23,29^. This is consistent with our conclusion that the transcriptome in seedling is oldest and most conserved at the amino acid sequence level, compared with root and flower because root and flower evolved later than stem and leaves.

By comparing their adaptive profiles, we obtained a global picture of the evolution of three important organs. Root, seedling and flower-specific sense-expressed genes are over-represented at the phase of Embryophyta, indicating an early adaptive peak of those organs at Embryophyta, although there is no fossil evidence supporting the origin of root, leaves and stems, and flower during this stage ^23,27,28,30^. Thus, the Embryophyta stage could have included preadaptive events for each organ, where genes that play a role in an organ initially emerged before the organ differentiated. Then from Magnoliophyta until Brassicales, organ-specific expressed genes show over-representation during these phylostrata, suggesting they are key stages for the evolution of these organs in the lineage leading to *A. thaliana*. A significant adaptive signal appearing in Magnoliophyta is expected, because fossil evidence suggests that innovation of real flowers originated within Magnoliophyta ^29^. Flowering plants are diverse as a group, with 250,000 to 400,000 species of Magnoliophyta ^31-33^. This compares to around 12,000 species of moss ^34^ or 11,000 species of pteridophytes ^35^. The adaptive peak of root and seedling at the stage of Magnoliophyta could result from the need for other organs like roots, stems, and leaves to adapt within that great diversity of Magnoliophyta species.

According to the Angiosperm Phylogeny Group III system (APGIII) ^36^ angiosperms (Magnoliophyta) can be divided into Eudicotyledons and Monocots. Between these, there are major differences in leaves, flowers, roots, and other organs ^37,38^, so it is meaningful that root, seedling and flower have an adaptive peak at this stage. Similarly, eudicotyledons, core eudicotyledons, Rosids and Brassicales include many different subgroups of plants. For example, core eudicotyledons include Rosids, Gunnerales, Dilleniaceae, Berberidopsidales, Santalales, Caryophyllales and asterids. These groups have large differences in their morphology, either in root, leaves or flower, which could explain the adaptive peaks in these phylostrata.

In higher eukaryotic organisms, 4% to 26% of protein coding genes are predicted to generate NAT ^39^. In *A. thaliana*, Yuan *et al*. identified 7,962 genes with antisense expression ^40^, somewhat higher than our results (6327). The difference could be due to the strict threshold (0.8 RPM, which is based on the distribution of the usually more-abundant sense expression) we set to define the genes with antisense expression. Our results showed that gene antisense expression tended to be organ-limited (expressed in either one or two of the three organs) compared to sense expression (Figure 1A & B; Table S1). NAT may played an important role in regulating the expression or translation of genes that are specifically expressed in a certain organ because 100% of the organ specific sense expressed genes have antisense expression (organ specific sense expressed genes: Root (1486), Seedling (191), Floral bud (1486)); in contrast, only 28.11% of the genes with sense expression shared by three organs (RSF: 16905) have antisense expression. This is supported by the result that genes with both sense and antisense expression have dramatically lower sense expression levels than genes with only sense expression (in seedling, root and flower, 29.8 vs. 58.0 RPM, P <0.001; 42.8 vs. 53.4 RPM, P <0.01; 34.9 vs. 52.5 RPM, P <0.001, respectively). However, patterns of antisense expression are qualitatively different from those of sense expression in important ways. Sense expression from all of our samples clustered strongly by organ, with only minor differences between accessions, but the antisense expression clustering into organs was weak and inconsistent (Figure 1C & D). Also, while TAI and TDI values clearly differed between organs in a relatively consistent pattern across accessions for sense expression, this was not true for antisense expression.

These major patterns of antisense expression call into question how much of the observed antisense expression is either functional or adaptive^41^. Another hypothesis compatible with our results assumes most of antisense expression is not adaptive (and indeed much may be maladaptive) but instead a noisy molecular byproduct of transcriptional machinery. Under this hypothesis, genes with the most critical function, highest expression, and/or widest expression might suppress this potentially maladaptive antisense byproduct, while other genes with lower or more limited expression and less critical functions may more frequently have antisense expression because negative selective pressure to eliminate their antisense expression byproduct is weaker. Though our data do not allow us to rule out either the adaptive or transcriptional noise hypotheses, a combination of both likely explains our observed antisense expression patterns.

The phylostratigraphic approach has been widely used to infer patterns of genome evolution, and has been applied to address questions related to the ontogenic patterns predicted by the developmental hourglass model, the adaptive history of different organs, and *de novo* gene origination ^9-12,16,17,42^. Central to this approach is the inference of the earliest emergence of target sequences at a particular phylogenetic node usually by using the similarity search algorithm BLAST ^43^ on a set of genomes that represent the nodes. BLAST has known limitations when sequences are highly diverged, especially in detecting remote homologues of short and fast-evolving sequences ^44,45^. Due to these limitations of the BLAST algorithm, published phylostratigraphic analyses have recently been criticized due to a predicted bias where certain short or fast-evolving genes will erroneously appear younger than they truly are as a result of BLAST false negatives. Through simulation, Moyers and Zhang argued that genomic phylostratigraphy underestimates gene age for 14% and 11% of the sequences they simulated for Drosophila and yeast, respectively ^45,46^. They argue that these potential errors for 11 or 14% of genes create spurious patterns in the distributions of certain gene properties, such as length or rate of evolution, across age groups. However, Domazet-Lošo, *et al*.^47^. re-assessed these simulations and identified problems that call into question the conclusions of Moyers and Zhang^45^. The problems include irreproducibility, statistical flaws, the use of unrealistic parameter values, and the reporting of only partial results from those simulations ^47^ Domazet-Lošo, *et al*.^47^ argued that, even with a possible overall BLAST false negative rate between 5-15%, the large majority (≥74%) of sequences assigned to a recent evolutionary origin by phylostratigraphy is unaffected by the technical concerns about BLAST^47^. Further, when they removed from the analysis genes that Moyers and Zhang^45^ found susceptible to the BLAST error in their simulations (192 out of 4157 genes with expression) the general profiles of biological patterns remained largely unaffected, indicating such potentially misplaced genes do not distort the major results^47^. They concluded that phylostratigraphic analyses of patterns of gene emergence and evolution are robust to the false negative rate of BLAST ^47^. Therefore, the phylostratigraphic approach we used has been proven to be valid, and there is no reason to suspect that our results are artifacts of our analytical procedures.

Taken together, the phylotranscriptomics approach, through combining gene expression and sequence evolution, works well to investigate the age and sequence divergence of the transcriptomes in different *Arabidopsis* organs and the adaptive profile of those organs along their evolutionary history. This study opens a new door in the investigation of plant organ evolution, and gives new indirect evidence on the adaptation of plant organs.

## MATERIALS AND METHODS

### Sample collection and transcriptome sequencing

Our study used seedlings (11-12 days old after 4 true leaves had emerged), root (from 10-day old seedlings) and floral bud (stage-12) RNA samples from the 19 *A. thaliana* accessions that are also MAGIC founders ^19,20^: Bur-0, Can-0, Col-0, Ct-1, Edi-0, Hi-0, Kn-0, Ler-0, Mt-0, No-0, Oy-0, Po-0, Rsch-4, Sf-2, Tsu-0, Wil-2, Ws-0, Wu-0 and Zu-0. Seedling and root tissue was collected from plants grown at 20^°^C under long day conditions (16 hours light 8 hours dark). In order to illicit nearly simultaneous flowering all accessions were vernalized for six weeks at 4^°^C under short day conditions followed by a shift back to long day conditions described above. Detailed procedures for plant growth, tissue collection, preparation of RNA, polyA-selected strand-specific library construction, and Illumina sequencing are as previously described ^19^. The RNA-seq read data for Col-0 and Can-0 were released previously as part of Gan *et al*.*^20^* under GEO accession numbers GSM764077-GSM764082. RNA-seq read data for the other 17 MAGIC founders have been released under GEO series GSE53197.

### Quantification of gene expression

The RNA-seq reads for all 19 MAGIC founders as available in the GSM764077-GSM764082 and GSE53197 releases for each of the three organs (seedling, root, and stage-12 floral bud) were aligned with PALMapper ^48^ to the *Arabidopsis* genome for read quantification after Gan *et al*.*^20^*. The quantification of gene expression and normalization were as described previously ^20^, and normalized expression (sense and antisense) of each gene for each organ from each accession (reads per million, or RPM) are available in Supplemental dataset1. For each gene, if its sense (or antisense) expression in a certain organ in at least 5 accessions was ≥ 0.8 RPM, then this gene was defined as sense (or antisense, respectively) expressed in this organ. We set 0.8 RPM as the threshold based on the distribution of the sense and antisense expression per gene per accession. In each organ half of the genes per accession have above 0.8 RPM sense expression, while less than 1% of genes per accession have above 0.8 RPM antisense expression.

In order to detect the overall similarity of sense and antisense transcriptome profiles across accessions and organs, we calculated Pearson’s correlation coefficient pairwise across all samples. Using the “heatmap.2” package in R, we represented the expression correlations in a heatmap, and performed clustering of samples using the Euclidean distance metric.

### Phylostratigraphy and *dN/dS* ratio

Our phylostratigraphic analysis was performed similarly to those presented previously ^9-11^. The data about each gene’s phylostratum were used directly from Drost *et al*. ^21^. Briefly, the first gene model for each *A. thaliana* gene from the TAIR10 release was assigned to one of 12 phylostrata, starting from the origin of cellular organisms and ending at *A. thaliana*. The age of each phylostratum, or the estimated time since the diversification of the corresponding clade from its most recent common ancestor, was extracted from Arendsee *et al*. ^49^.

The data on the dN/dS ratios for each gene were used directly from Quint, *et al*. ^11^. Specifically, orthologous gene pairs of *A. thaliana* and *A. lyrata* or the closely related Brassicaceaes, *T. halophila, C. rubella* or *B. rapa*, were determined with the method of best hits using blastp ^50-54^. And dN/dS ratios of only those orthologous gene pairs with dN<0.5, *dS*<*5* and *dN/dS* < 2 were retained for further analysis ^11^.

### Calculation of TAI and TDI

As described previously ^10-12,21^, the TAI of organ s was calculated as the weighted mean of the evolutionary age (PS)*ps_i_* of gene *i* weighted by the expression level *e_is_* of gene *i* at organ *s*:

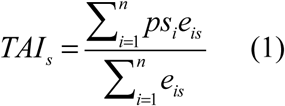
where *n* is the total number of genes analyzed. And the expression level was estimated as RPKM. Low PS values correspond to evolutionarily old genes, thus the lower the TAI value, the older the transcriptome is. Likewise, high PS values correspond to evolutionarily young genes, so the higher the TAI value, the evolutionarily younger the transcriptome is.

Analogously, the TDI of organ *s* was calculated as the weighted mean of the sequence divergence (*dN/dS*) of gene *i* weighted by the expression level *e_is_* of gene *i* at organ *s*
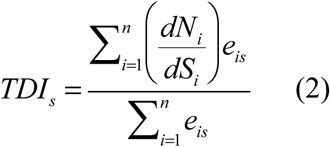
where *n* is the total number of genes analyzed. Low/high *dN/dS* ratios correspond to conserved/divergent genes, so low/high TDI values correspond to conserved/divergent transcriptomes. To investigate the dependence of the results on the reference species for the calculation of TDI, the analysis was repeated for each reference species (*T. halophila, C. rubella* and *B. rapa*). The results are presented in Figure S2 and Tables S6-11.

### Estimating standard errors of TAI and TDI by bootstrap analysis

The TAI is written alternatively as the sum of products between the phylostratum *ps_i_* of gene *i* and the partial concentration *f_is_* of gene *i* at organ *s*
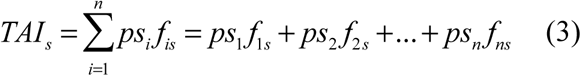

Analogously, the TDI is written alternatively as:
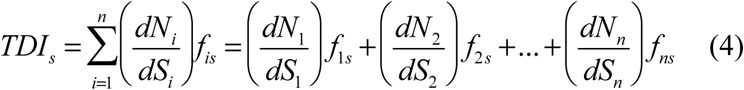
where the partial concentration *f_is_* is calculated as the expression level *e_is_* from equation (1) divided by the denominator of equation (1) and *n* is the total number of genes analyzed.

Bootstrapping was used to approximate the standard errors of TAI (or TDI) ^11,12^. In brief, for each accession, resampling 25,078 genes (25,078 out of 25,260 genes with an assigned phylostratum have expression) with replacement was done 1,000 times. Then, using the phylostratum, dN/dS ratio, and expression abundance (RPKM) in each organ associated with each gene, TAI and TDI can be computed according to equations (3) and (4) for each bootstrap sample of genes. A sample mean of TAI and TDI was obtained according to the results of the 1,000 bootstrap samples. Similarly, the standard errors of the distribution of the sample means were calculated based on the results of the 1,000 bootstrap samples, and were used to estimate the standard error of TAI (or TDI).

### Statistical significance of the TAI and TDI profiles

For each accession, we performed the Mann Whitney test on TAI and TDI for comparison of each pair of organs based on the results obtained from 1,000 bootstraps.

### Relative expression of genes for a given phylostratum

As described previously ^10-12,21^, the relative expression *RE_ls_* of the genes in PS *l* in organ s was calculated as
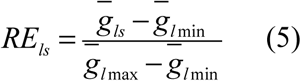
where 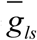 denotes the average partial expression of the genes in PS *l* at organ s and 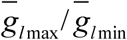 is the maximum/minimum average partial expression across all organs. The value of *RE_ls_* ranges from 0 to 1, where 0 denotes the organ where genes in PS *l* show minimum average partial expression and 1 denotes the organ where genes in PS *l* show maximum average partial expression. We further grouped the relative expression levels into two PS classes, where the first PS class consists of relative expression levels of genes belonging to the three oldest phylostrata PS1-PS3 and the second PS class consists of relative expression levels of genes belonging to the younger phylostrata PS4-PS12^11^. This grouping was chosen because it distinguishes phylostrata unique to land plants (PS4-PS12) that are evolutionarily relatively young, from older phylostrata (PS1-PS3) that arose before the origin of land plants.

### Phylostrata enrichment calculations for organ-specific expressed genes

If organ-specific expressed genes are enriched in a certain phylostratum, this indicates that there are adaptive signals in this phylostratum ^16^. In order to find the adaptive signals of three main organs of *A. thaliana*, we firstly defined the organ-specific expressed genes (indicated as R, S and F), separately for sense and antisense expression, and then mapped those genes back to the phylostratigraphy profile that spanned 12 phylostrata ^11,16,17,21,55^. In order to have another set for comparison, we also mapped the genes expressed in three organs (RSF) to each phylostrata.

For each of the groups R, S, F and RSF, we performed an over-representation analysis by comparing the actual frequency of each group in a phylostratum with their expected frequency based on the product of the marginal frequencies of the four groups and the proportion of all genes with assigned phylostrata found within each of the 12 different phylostrata ^16,17^. The logarithm of the ratio of actual and expected frequencies, or log-odds ratio, was obtained to indicate the enrichment of a group within a certain phylostratum. Then we performed the two-tailed hypergeometric test ^56^ to test the significance of the over- or under-representation, and controlled for multiple comparisons through a Bonferroni correction.

### Gene Ontology for organ-specific expressed genes

Enrichment analysis of gene ontology (GO) terms was performed on organ-specific sense-expressed genes (R, S and F) and genes sense-expressed in all three organs (RSF) using Agri GO ^57^. All GO analyses used as background all genes with sense expression in at least one organ. The significance was assessed with hypergeometric tests and corrected by False Discovery Rate. The significance threshold was set at P=0.05, and GO annotations that did not appear in at least 5 entries were not shown, that is, the minimum number of mapping entries was set to 5.

## Acknowledgments

We thank Dr. Richard M. Clark for comments on the manuscript. We also thank Dr. Rätsch Gunnar for his contribution to processing the RNA-seq data and getting the gene expression estimates. This work was supported by the National Science Foundation (NSF) (0929262 to C. T. and Richard M. Clark) and Kansas Institutional Development Award (IDeA) Network of Biomedical Research Excellence (K-INBRE) Postdoc Award (to L.L) from the National Institute of General Medical Sciences of the National Institutes of Health (NIH) under grant number P20 GM103418. Contribution number 16-246-J from the Kansas Agricultural Experiment Station.

## Contributions

L.L. and C.T. conceived the study, managed its organization and edited the draft versions of the manuscript. L.L. performed research (all the different steps), analyzed data and wrote the first draft of the manuscript. E. J. O. performed research (expression data analysis), J. G. S. performed research (producing the RNA-seq data). L.L., E. J. O., J. G. S. and C.T edited the manuscript. All authors discussed the results, commented on the manuscript, read and approved the final manuscript.

